# Robust higher-order multiplexing in digital PCR by color-combination

**DOI:** 10.1101/2023.05.10.540190

**Authors:** Irene Santos-Barriopedro, Sylvain Ursuegui, Etienne Fradet, Rémi Dangla

## Abstract

There is a growing need in molecular biology to interrogate samples for higher number of biomarkers, beyond the 2 to 5 biomarkers typically addressable with standard PCR technologies. Here, we demonstrate a novel approach to increase the level of multiplexing in digital PCR up to 15-plex by detecting each target with 2 distinct fluorophores with a 6-color digital PCR system, a method called digital PCR by color combination. We provide a statistical framework to interpret digital PCR data by color combination, predicting that high-plexed assays by color combination can, in theory, have the same precision and sensitivity as corresponding single-plex assays. We verified the statistical framework and confirmed the sensitivity prediction experimentally on a 15-plex assay and an 11-plex assay. These results establish the technical framework for future high-plex assays addressing a variety of applications and utilities, such as infectious disease syndromic panels or tumor mutation profiling panels.

## Introduction

In the last decade, the knowledge of genome sequence landscapes has been revolutionized. Tremendous advances have been made in the understanding of the differences and similarities among genomes within a species and across species since the completion of the Human Genome Project (HGP) in 2003. For example, we now know that the genomes of all individuals are nearly identical, and variations, such as a single nucleotide changes, in approximately 0.1% of the genome sequence account for many of the differences among us. Determining these differences in the genome is critical to understand the principles and mechanisms of human health and disease and tracking emerging sequence differences can facilitate disease prognosis and elucidate the efficacy of a treatment. Indeed, the identification of a growing set of genetic biomarkers and genetic signatures associated with a range of diseases has enabled the implementation of cost effective and sensitive techniques to follow their evolution over time and throughout treatment cycles in blood and other body fluids (liquid biopsies), and tissues.

One such technique, digital PCR (dPCR), has seen rapid adoption since the availability of commercial instrumentation as it is capable of single molecule detection sensitivity and precise quantitation of targets, even in complex samples such as liquid biopsies^1,2^. In addition, it offers a practical solution to the shortcomings of real-time quantitative PCR (qPCR) including improved sensitivity, precision, and resistance to inhibitors while leveraging similar PCR chemistry and workflow.

Until now, instrumentation and assays for qPCR and dPCR have been typically limited to the characterization of only a handful of nucleic acid targets in a single sample, i.e., low-plex assays. With the growing demand across multiple fields of molecular biology for mid- to high-plexed assays that interrogate tens to hundreds of targets in parallel, thereby giving a more thorough genetic characterization of the interrogated sample, the need for high-plex assays is more critical than ever. Such high-plex assays are accessible through advanced technologies, such as microarrays and next-generation sequencing (NGS), but these technical solutions come at the expense of complex and multi-day workflows and high costs per sample. Thus, by increasing the plexing capacity of a single PCR assay while maintaining the simplicity and high-performance workflow, dPCR stands to bring high added value to nucleic acid detection workflows across multiple applications.

The most simplistic approach to expand the multiplexing capability of qPCR or dPCR is to increase the number of fluorescent detection channels, or colors, while using each color to detect one nucleic acid target. With this “1 Color = 1 Target” approach (1C-1T multiplexing) (Figure 1A), a maximum of a 3-plex can be achieved with a 3-color detection system; or a maximum of a 6-plex with a 6-color detection system. As assay design and data analysis is relatively straightforward, this approach has motivated qPCR and dPCR instrument providers to increase progressively the number of detection channels accessible across their instrumentation^3,4^. However, the number of colors cannot be increased infinitely due to the limited bandwidth of optical receptors and the spectrum overlap of commonly available fluorescent reporters.

**Figure 1.**
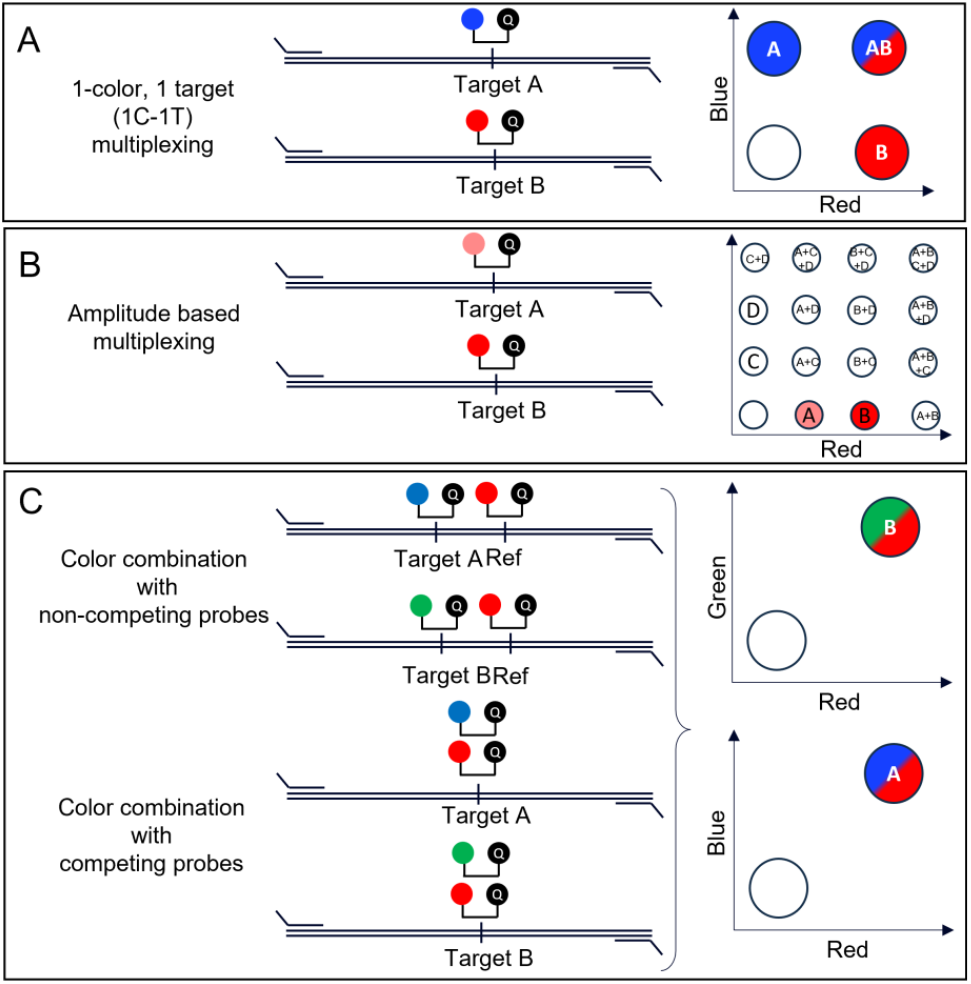
Schematics of different multiplexing strategies: **(A)** The 1C-1T multiplexing approach. Each target is detected using a single TaqMan® probe, each labeled with a unique fluorophore. For the blue/red duplex assay shown, four clusters will be visible in the 2D dotplot: all negative partitions, blue for Target A only positive partitions, red for Target B only positive partitions and blue/red for Target A & Target B positive partitions. **(B)** The “amplitude-based” multiplexing approach. Two distinct targets are detected using the red channel, one with a low fluorescence intensity cluster (Target A) and one with a high fluorescence intensity cluster (Target B). Similarly, two targets (C and D) are shown as detected using a second color. Using this approach with two colors leads to up to 16 clusters in the 2D dot plot (accounting for double, triple and quadruple positive cluster combinations in addition to the single positive and negative clusters). **(C)** The two “color combination” approach, with 2 different implementations that lead to a similar fluorescence data structure. The cluster of droplets that contains Target A only is blue/red while the cluster of droplets that contains Target B is green/red. Q represents the Fluorescence Quencher molecule conjugated to the probe.

Consequently, more sophisticated approaches have been implemented to increase the level of plexing beyond that offered by 1C-1T multiplexing. These approaches are commonly referred to as *high-order multiplexing methods* ^5,6^. A first concept, called *amplitude-based* or *intensity-based* multiplexing, consists of detecting multiple targets within the same color channel by varying probe concentrations and, by consequence, their emitted fluorescence intensities within the same color channel^7–9^ (Figure 1B). A similar concept, called *ratio-based multiplexing*, consists of detecting targets by varying the ratio of 2 probes labeled with fluorophores emitting in different color channels of each target^10,11^.

A shared drawback of the *high-order multiplexing* methods referenced above is an increase in the complexity of the assay design and data analysis, as well as a loss in assay robustness. For example, when implementing *amplitude-based* multiplexing in dPCR, at least 3 different intensity thresholds must be set for each detection channel to properly classify the partitions according to their fluorescence intensity. Frequently, thresholding is impractical, and each individual detected cluster must be identified on 2D dot-plots, requiring 16 clusters to be identified for a simple 2-color 4-plex assay. In addition, variations in fluorescence intensities from sample-to-sample lead to high risks for partition misclassification with increased rates of false-positive and false-negative calls.

Alternative approaches to increase the multiplexing capabilities of qPCR have been proposed around the concepts of *multicolor combinatorial probe coding*^12–14^ or *color-coding*^15^. In these approaches, each target is encoded using a unique combination of two or more distinct fluorophores, or colors. When two colors are used for each target, the maximum level of multiplexing with *n* colors is 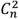 Using the color-coding method, Marras et. al.^15^ demonstrated a 15-plex assay using a 6-color qPCR instrument for the detection, identification and quantification of 15 different bacterial species.

Although certainly innovative, these qPCR methods have two major drawbacks. The first is that they are semi-quantitative. The second is that they work only if a single target is present in the sample. When more than one target is present, the method is ambiguous and incapable of reliably identifying the present targets. For example, in a colorcoded qPCR assay set up using the three colors, blue/green/red if a sample turns out positive for all three colors, it could be due to the simultaneous presence of either the blue/green and green/red targets or the blue/green and blue/red targets. Marras et. al. hinted to the potential for digital PCR to address these limitations in the qPCR workflow due to the unique sample partitioning step of digital PCR.

In this study, we implemented high-order multiplexing in digital PCR by color combination using a set of unique color pairs per target. Compared to amplitude-based or ratio-based approaches, the data analysis is simplified to a binary classification of each color, enhancing assay robustness. We studied the underlying statistical framework to evaluate the analytical power of the method and its limitations. We leveraged the 6-colors naica® system from Stilla Technologies to prepare a 15-plex and a 11-plex demonstration assay and verified the statistical predictions, in particular the sensitivity of the method.

## Results

### High-order multiplexing in digital PCR by color combination

We describe two possible strategies for color combination in digital PCR.

In the first dPCR strategy, two TaqMan® probes with two different colors are combined for each target but each TaqMan probe binds to adjacent sequences such that they do not compete to hybridize during PCR (Figure 1C - Color Combination with non-competing probes, upper scheme). In the illustrated example,, one probe targets the specific sequence (blue and green) while a second probe targets an adjacent reference sequence (red). During the PCR reaction, fluorophores of both colors are cleaved by the polymerase at each cycle.

In the second dPCR strategy, two TaqMan probes with two different colors are designed to hybridize to the same target sequence during the PCR reaction. Consequently, they compete to hybridize to the same binding site (Figure 1C, Color combination with competing probes, lower scheme). Here, during the PCR reaction, only one fluorophore is cleaved by the polymerase from each single-stranded DNA at each cycle but, if both probes have a 1:1 competition ratio, the same amount of fluorophore is cleaved on average for each color at the end of the PCR reaction.

Both strategies produce a similar fluorescence data structure in which partitions can be clustered depending on a simple binary “negative” versus “positive” call for each color used in the assay, similar to the classical 1C-1T multiplexing. Hence, the straightforward and robust 1D or 2D thresholding methods to analyze 1C-1T data are applicable to high-plexed color combination data, in contrast to high-plexed *amplitude-based* data, which requires more complex thresholding methods.

Thus, high-plexed dPCR by color combination can be viewed in a manner similar to classical 1C-1T dPCR, except that a partition containing one target will be positive for at least two colors at the same time.

### Statistical considerations for color combination

In dPCR, the targeted template molecules initially present in the reaction volume are randomly distributed across a large number *N*_*tot*_ of partitions and subsequently amplified by PCR. The partitions are then read by a fluorescence imager and classified as negative (i.e., do not contain the targeted template molecule *T*), or positive (i.e., do contain at least one targeted template molecule *T*). The statistical framework to convert the count of negative and positive partitions into an estimation of the concentration C_*T*_ of the target *T* relies on the Poisson approximation and has been described previously^16^ for 1C-1T dPCR:

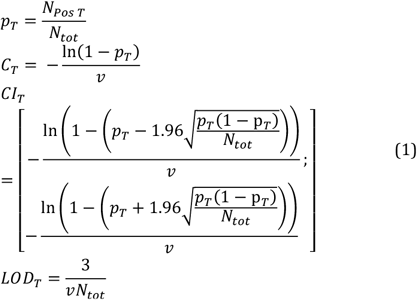

where *p*_*T*_ is the estimator for the probability of a partition to contain at least one targeted template molecule *T, N*_*Pos T*_ is the number of partitions classified as positive for the target *T, v* is the average volume of the partitions, *CI*_*T*_ is the 95% confidence interval for C_*T*_ and *LOD*_*T*_ is the theoretical concentration at limit of detection for the target *T*.

For a multiplexed assay targeting *i* distinct targets *T*_*i*_, the same formulas apply to each target if all targets are statistically independent (i.e., on different template molecules or fragments), and as long as the target content of all partitions can be unambiguously inferred from the fluorescence data.

The latter constraint cannot be satisfied when higher-order multiplexing by color combination is implemented because some clusters of partitions are ambiguous regarding their target content. These ambiguous clusters are those that are positive for strictly more than two colors. For example, in the case of a 3-plex assay set up with three colors (blue/green/red), there is a total of eight possible classes of partitions when classified according to the target contained in a partition (Figure 2A).

**Figure 2.**
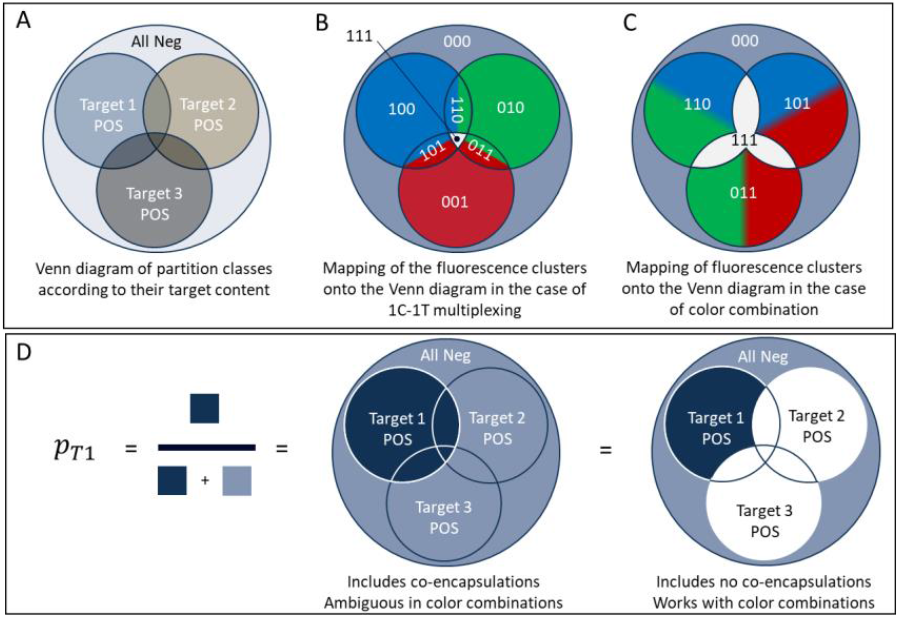
Representations of cluster possibilities following different multiplexing strategies. **(A)** Venn diagram of the 8 possible classes of partitions, classified according to their target content. **(B)** Overlay of the fluorescence clusters on top of the Venn diagram in the case of a 3-plex assay set up using 1C-1T; the color of a clusters corresponds to the detection channel for which the partitions are positive in this cluster; the triple positive cluster 111 is white. **(C)** Same overlay of fluorescence clusters but in the case of a 3-plex assay set up using color combination. The triple positive cluster is white. D) Color coded Venn diagram of the 8 possible classes of partitions showing groups of partitions used in the calculation of 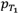

When using 1C-1T, the droplets can be grouped into eight clusters depending on their fluorescence (Figure 2.B). For nomenclature, we name 000 the *fluorescence cluster* of droplets that are negative for all fluorescence channels, 100 the cluster of droplets positive in blue but negative in green and red, 010 the cluster of droplets negative in blue, positive in green, and negative in red, etc… for 1C-1T, there is a direct correspondence between the *target content class* and the *fluorescence clusters*, such that there is no ambiguity to extract target content from fluorescence clusters (Figure 2A and 2B).

However, when using color combination with two different colors per target, there are only five possible fluorescence clusters (Figure 2C). The 111 fluorescence cluster maps to the union of four target content classes: a droplet in the cluster 111 could contain Target 1 and Target 2, or Target 1 and Target 3, or Target 2 and Target 3, or Target 1 and Target 2 and Target 3. In other words, a droplet in the 111 cluster can contain either the blue/green target and the blue/red target, or the blue/green target and the green/red target, or the blue/red target and the green/red target, or all three targets together. Therefore, there is no way to infer the content of a partition that is blue/green/red positive. Thus, the triple positive cluster 111 is ambiguous with regards to its target content.

This ambiguity holds true for all clusters in which partitions are positive for more than two of the colors used in color combination. When implementing color combination with six detection channels, clusters that are triple positive, quadruple positive, etc… are all ambiguous.

Owing to this ambiguity, the set of equations (1) applied to 1C-1T data cannot be applied to color combination data. One mathematical solution consists of discarding all ambiguous clusters of positive partitions that may contain a given target *T*_*i*_. To be statistically consistent and obtain an accurate estimation of 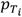 the clusters of partitions that are negative for target *T*_*i*_ but positive for another target present in at least one of the discarded ambiguous clusters must be discarded as well from the total number of partitions included in the estimation of 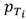 This rational is visually explained in Figure 2D.

Consequently, when using color combination, an accurate estimation of the concentration 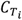of a given target *T*_*i*_ is obtained using the set of equations:

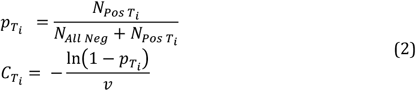

where 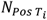 is the number of partitions in the cluster of partitions that are unambiguously positive for target *T*_*i*_ only (and negative for all other targets) and *N*_*All Neg*_ is the number of partitions negative for all targets.

When using two colors to detect each target, the partitions that are positive for a given target *T*_*i*_ are in the cluster of partitions that are positive for only the two colors corresponding to target *T*_*i*_. And the partitions that are all negative are in the clusters of partitions negative for all colors. In theory, if color combination is used to detect all targets in the assay, there should not be any cluster of partitions positive for a single color. In practice, there may be a few such partitions due to noise in the fluorescence signals of the assay. These partitions can be either included as all negative partitions or discarded from the analysis.

The exclusion of a set of partitions in the detection of target *T*_*i*_ and the estimation of *C*_*i*_ leads to a loss in sensitivity and precision. The formulas for the 95% confidence interval and the LOD for *C*_*i*_ are (see Supp. Mat. #1 for calculations):

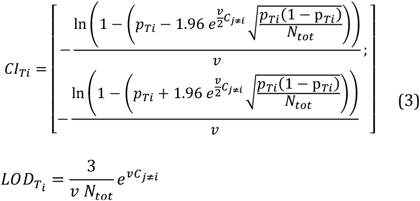

Both formulas include an exponential factor that involves 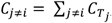 which is the sum of the concentrations of the targets other than *T*_*i*_ detected in the color combination assay, i.e. the background concentration of targets other than *T*_*i*_.

Interestingly, the LOD with color combination 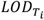scales as the LOD for the 1C-1T approach *LOD*_*T*_ times the mentioned exponential factor, which can be highlighted by defining the LOD contrast Δ*LOD* (in %):

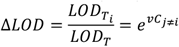

Figure 3A shows a plot of Δ*LOD* against the background concentration for typical sizes of droplet partitions. For the size of droplets produced on the Sapphire Chip and Ruby Chip of the naica® system, the increase and degradation in LOD when using color combination is less than 20% for background concentrations below 300 copies/μL. However, a high-order color combination assay also brings a net gain in sensitivity when compared to an equivalent 1C-1T panel of assays by avoiding the splitting of the sample across multiple different reactions. For example, consider a 15-plex assay performed either with the color combination or with 1C-1T multiplexing, both read out on a 6-channel system. The 15-plex assay can be performed in a single well with color combination but would require being split across three different wells using the 1C-1T strategy. As a result, the LOD of the 15-plex assay using 1C-1T will be three times higher than that of the 15-plex using color combination. In this example, the LOD contrast becomes:

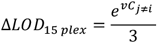

**Figure 3.**
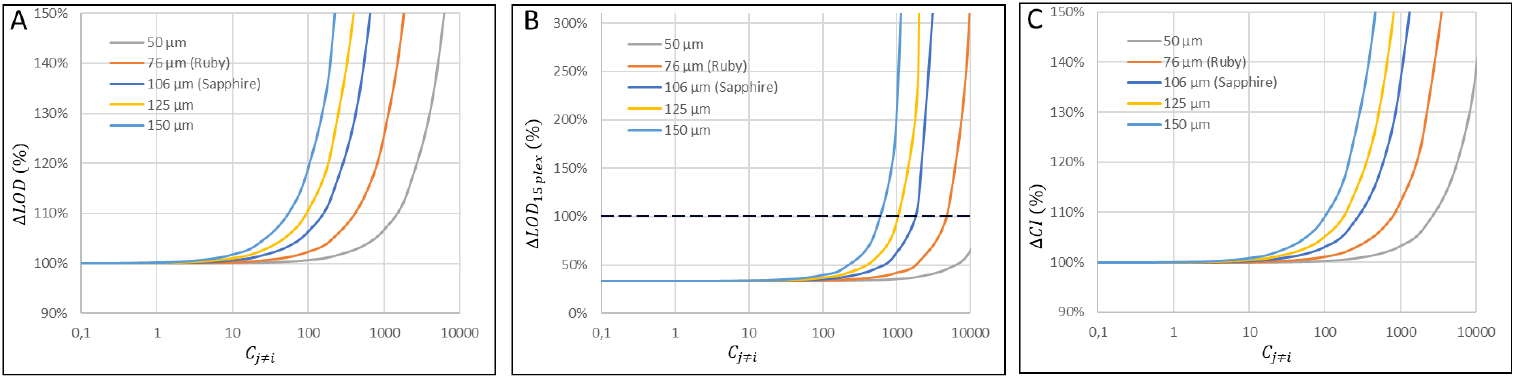
Plots of. ***(A)*** *ΔLOD* in %, ***(B)*** *ΔLOD*_15 *plex*_ in % and ***(C)*** *ΔCI* in % versus the background concentration *C*_*j*≠*i*_ in copies/μL for five different partition diameters and volumes, including the partition volumes of the Sapphire Chip and the Ruby Chip of Stilla Technologies.

Thus, the LOD of the 15-plex color-combination assay is consistently superior to the LOD of an equivalent 1C-1T panel of assays detecting 15 targets for a background concentration of up to more than 1000 copies/μL (Figure 3B). Similarly, the width of the confidence interval (CI) for color combination scales as 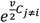 (Figure 3C). The relative degradation in precision is below 20% for background concentrations below 1000 copies/μL for the volume of partitions in a Ruby chip.

It is possible to keep the analysis framework described above for a 2-color color combination assay and add one more target *T*_*L*_ to the assay by targeting it with a single color. For example, using the three colors blue/green/red, three targets can be detected by color combination as blue/green, blue/red and green/red, and one additional target can be detected as blue-only. In this case, the blue/green and blue/red targets require an updated equation to account that some blue/green and blue/red droplets can also contain the blue-only target.

The generalized formulas for this configuration are:

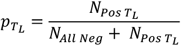

and, if one of the colors used to detect target *T*_*i*_ is also used as a single color to detect the last target *T*_*L*_, then

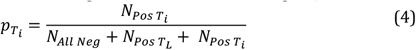

Otherwise

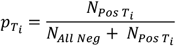

where 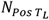 is the number of partitions that are positive for the single color of *T*_*L*_ and 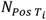 is the number of partitions that are positive for only the two colors used to detect target *T*_*i*_.

In applications such as liquid biopsy analysis for which having the best sensitivity is preferable, the loss of sensitivity due to the background concentration may be problematic. In the case of liquid biopsy, the highest contributor to the background concentration is most likely to be the target used to quantify the wild-type (WT) circulating free DNA (cfDNA). By using a single color to detect the WT target and using color combination with the other available colors to detect the low concentration mutant targets, optimal assay sensitivity can be obtained.

Using this color combination with one single color approach with a 6-color digital PCR system, it is then possible to setup a 12-plex assay composed of one wild-type target + 11 mutant targets. Here, one color is allocated to the detection of the wild-type target and the five additional colors are combined pairwise to detect 10 mutant targets with one additional mutant target also detected using one of the five additional colors.

With this assay configuration, the set of equations (1) can be used for the WT target as it is unambiguous. For the mutant targets, the set of equations (2) must be slightly modified to add the WT positive droplets to the total analyzed droplets and thereby improve the sensitivity:

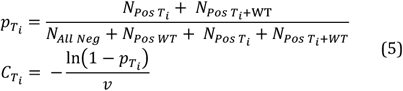

Here, the concentration of the WT target isolated in one color has no impact on the sensitivity and precision of the measurements of the mutant targets.

### Demonstration of a 15-plex assay with color combinations

To verify the feasibility of color combination in dPCR and test the predictions from equations (2) and (3), a proof-of-concept 15-plex dPCR assay was developed (Table 1 and detailed in the Supp. Mat. #2, #3 & #7).

**Table 1:**
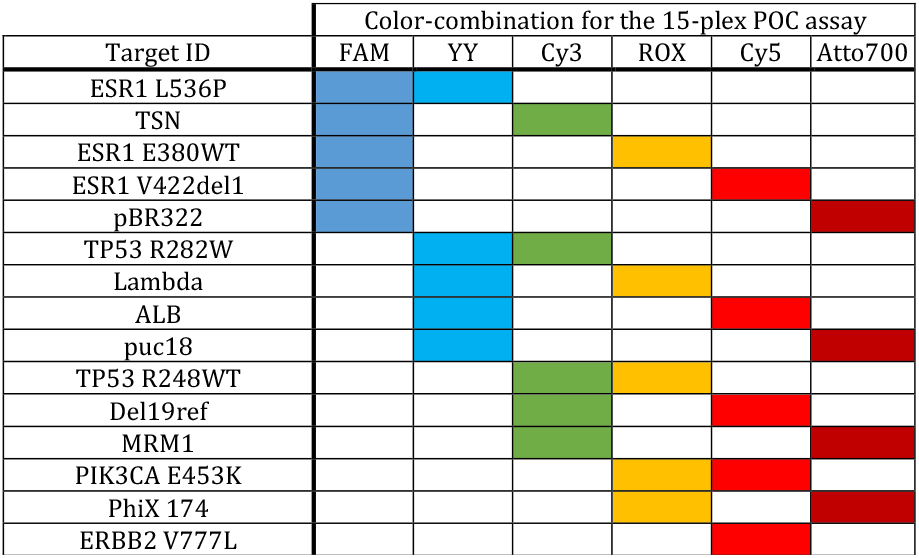
Structure of 15-plex proof-of-concept assay. 14 targets are detected using 2 colors each, as indicated by the colored cells of the table. The last target is detected using only a Cy5 probe.

Using gBlocks™ gene fragments (Integrated DNA Technologies, Coralville, Iowa, USA), the concentration of each target was adjusted individually to test the theoretical framework developed herein. Figures 4A and 4B show an example of the data obtained with this assay, analyzed using the 6-color naica® system (Stilla Technologies, Villejuif, France).

**Figure 4:**
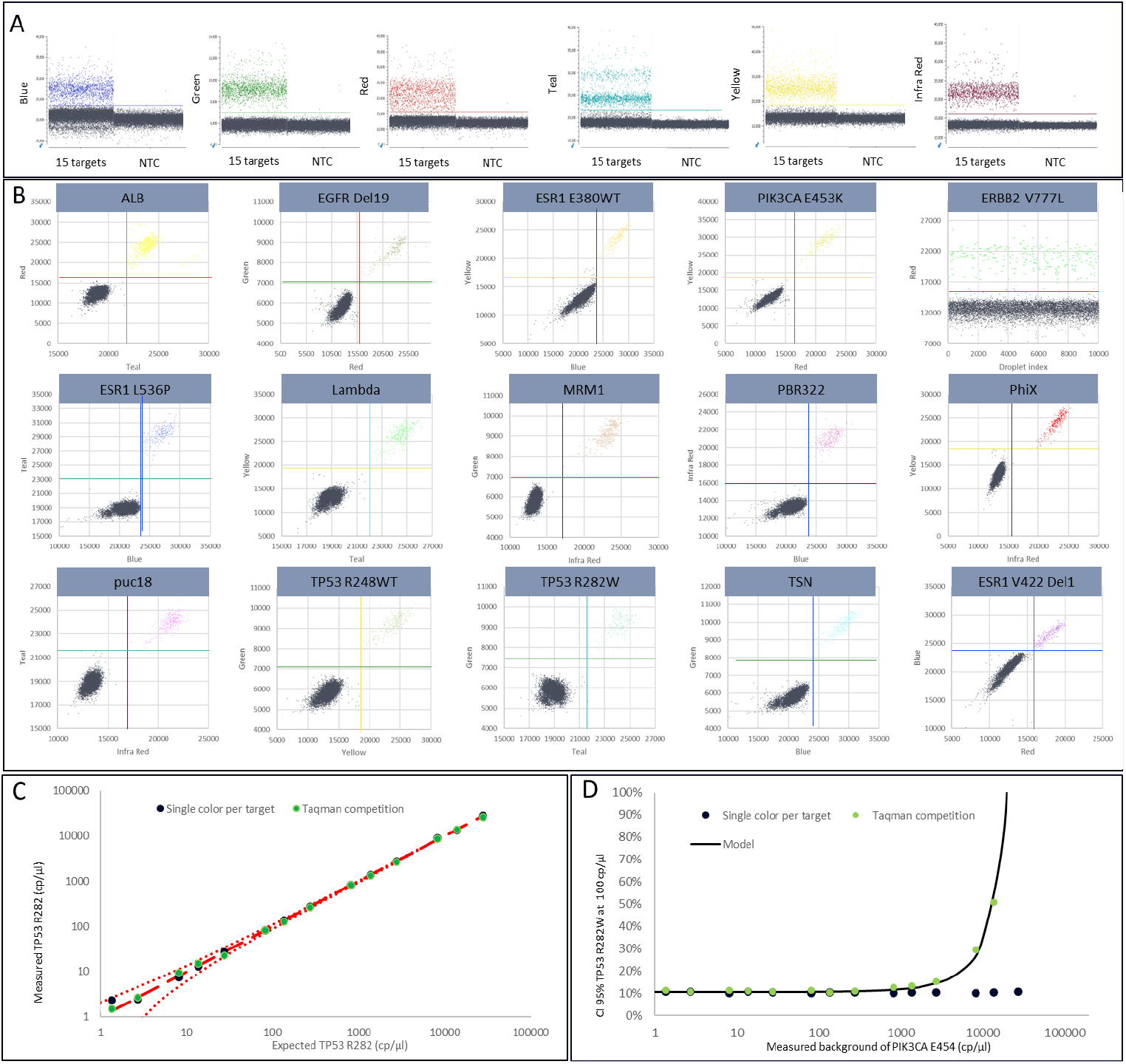
**(A)** The six 1D dot-plots obtained for the 15-plex POC assay with all target concentrations at 100 copies/μL using the Crystal Miner software. Relative fluorescence intensities corresponding to each color channel are plotted on the Y-axis while the assay DNA template used is specified on the X-axis. NTC denotes the no template control. **(B)** The 14 truncated 2D dot-plots, each one corresponding to a combination of two colors used to detect one target, and the 1D truncated dot-plot for the detection of the Target ERBB2 V777L using Cy5 only. The data is truncated in a way that, for each 2D dot-plot, the data that is above the positivity threshold for the other four non-visible dimensions are not projected onto the 2D dot-plot but discarded from the visualization. This visualization corresponds to the view of the partitions that are included in the set of equations (2). For the 2D dot-plots, relative fluorescence intensities corresponding to each color channel are plotted on the X- and Y-axes. For the 1D dot-plot, the relative fluorescence intensity corresponding to the red color channel is plotted on the Y-axis and the position of each droplet corresponding to the droplet index is plotted on the X-axis. **(C)** Plot of the measured vs expected concentration for Target R282W, as measured with single color per target (1C-1T) or with color combination using competing TaqMan® probes; the red dotted lines show the width of the predicted 95% confidence interval. A background of 100 copies/μL of PhiX DNA was added to the color combination and a background of 100 copies/μL of ALB DNA was added to the 1C-1T configuration. **(D)** Plot of the measurement confidence interval, as provided by the Crystal Miner software (Stilla Technologies), for Target TP53 R282W at approximated 100 copies/μL and measured in a varying background concentration, both with single color per target (1C-1T) and with color combination competing TaqMan® probes; the black line is the prediction from equation (3).

In this example, all targets are present at concentrations of approximately 100 copies/μL each in the Ruby Chip chamber. Yet, in contrast to the case of color-coded qPCR, each target was correctly detected and precisely quantified. Binary classification of the partitions for each color was achieved by placing six linear thresholds on 1D dot-plot of each color (Figure 4.A) and the statistical framework described above to calculate the concentrations of each target was applied automatically by the Crystal Miner software (Stilla Technologies) using the population editor tool (see Suppl. Mat. #8) It is noteworthy that in this assay configuration, the human eye is incapable of extracting information from the 1D dot-plots that are projections of 6-dimensional data with multiple double-positive, triple-positive etc… clusters.

The 14 truncated 2D dot-plots (Figure 4B), each display the cluster of double positive partitions for one target. In this view, the 1D thresholds combine correctly for each target to separate the all-negative cluster from the double positive cluster. Untruncated dot plots are shown in Supp. Mat. #4. To verify the concordance of the concentration measurements obtained with the set of equations (2) compared to those of set (1), a dilution series of the Target TP53 R282W was performed in a background of 100 copies/μL for one target in two configurations: i) with the 15-plex color combination assay and ii) with a comparable 1C-1T assay (see supplementary information for a description of the 1C-1T assay). All measurements were concordant across the tested dynamic range (Figure 4C).

Similarly, the prediction (3) for the confidence interval was evaluated with the same experiment. The experimental measurements of the confidence interval match with the prediction from equation (3) (Figure 4D).

Finally, the prediction (3) for the loss in sensitivity and limit of detection was tested by measuring the LOD of the Target *TP53 R282W* with a background concentration of the other targets at 100 copies/μL with and without using color combination. The LOD without color combination was measured at 1.02 copies/μL in the Ruby Chip chamber while the LOD with color combination was measure at 1.34 copies/μLin the Ruby Chip chamber, a value similar to the prediction from equation (3) of 1.42 copies/μL.

### Demonstration of 10+1-plex proof-of-concept

As described above, the detection of a possibly highly concentrated wild-type target on a single color theoretically overcomes the problem of loss of sensitivity in color combination. To demonstrate experimentally the prediction (5), a proof-of-concept 11-plex assay was setup (Table 2). In this experiment, 10 genes were each detected with two colors and one target (*PIK3CA E453K*) was detected with the single-color red. Similar experimental conditions as in the 15-plex assay were used (see Supp. Mat. #5 & #7). This assay was also compared to an equivalent 1C-1T assay containing the same 11 targets (see Supp. Mat. #6).

**Table 2:**
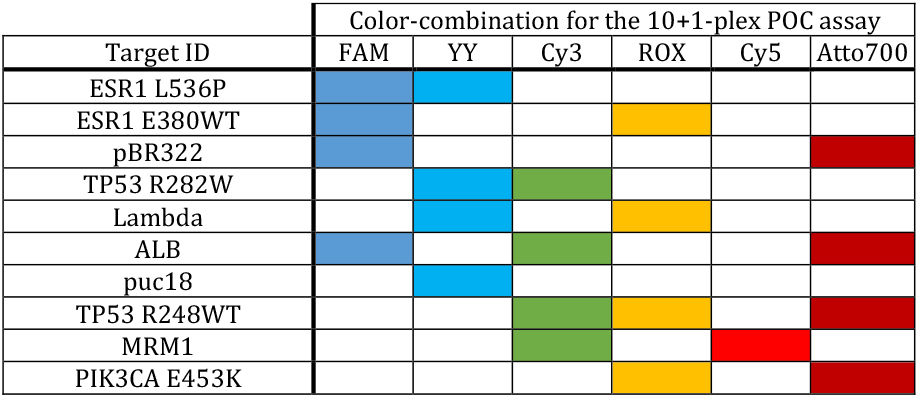
Structure of 10+1-plex POC assay where 10 genes are assigned to two colors and one gene is assigned to a different one single color.

Both the expected versus measured concentrations of TP53 R282W for 11-plex assay and the concentrations measured by the color combination assay versus the equivalent 1C-1T assay showed strong concordance across the entire tested dynamic range (Figure 5A). In contrast to the 15-plex assay configuration, the confidence interval remained unperturbed for a gene with color combination (*TP53 R282W*) when the concentration of the single-color gene (*PIK3CA E453K*) was increased (Figure 5.B) because droplets containing the single-color gene were not discarded from the analysis.

**Figure 5.**
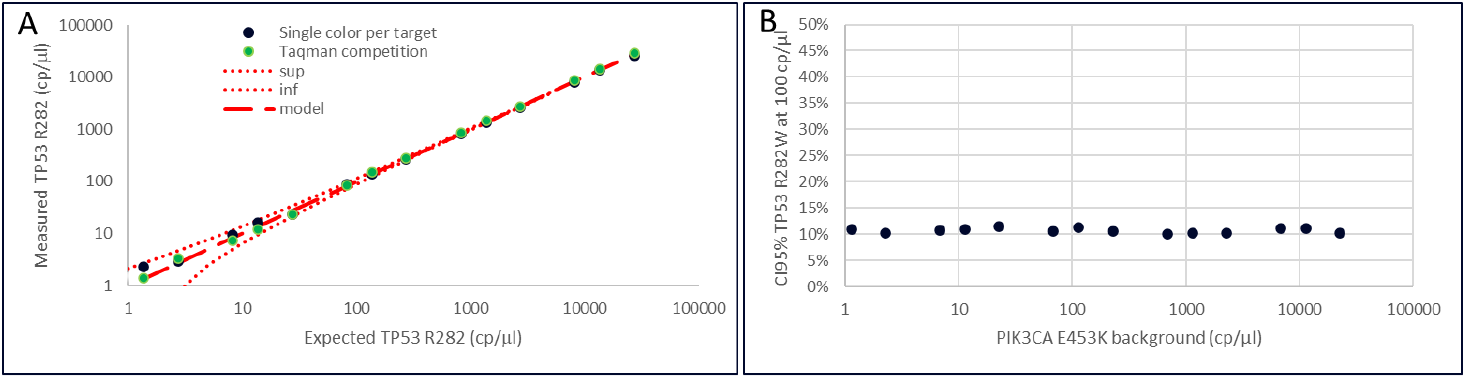
– 10+1-plex assay: 10 genes are attributed to two colors and one gene is detected by one single color different from the 10 genes colors. ***(A)*** Different concentrations of TP53 R282W measured in a 1000 cps/μl background of PIK3CA E453K. TP53 R282W measured concentrations are plotted against the expected concentrations. Experiments have been performed with color combination setup (table 2, TaqMan competition) and with one single color per target (details in Supp. Mat.). ***(B)*** CI 95% graphics. CI95% obtained for varying PIK3CA E453K concentrations with a background of TP53 R282W set at 100 cps/μl.

Lastly, the LOD of the Target *TP53 R282W* in this 11-plex assay configuration was measured by both color combination and 1C-1T. The background gene was a single-color gene (*PIK3CA E453K*) for which the concentration was set at 1000 cp/μl. No obvious difference in LOD was observed with measurements of 1.22 cp/μl with color combination and 1.21 cp/μl using 1T-1C.

In summary, this confirms that allocating one color to the wild-type target and combining the other colors for mutant detection allows to accurately quantify all targets with LODs similar to 1C-1T.

## Discussion & Conclusion

Digital PCR is an affordable and sensitive alternative to NGS for the monitoring of rare events but still lacks the multiplexing power that NGS offers. Our results show that it is possible to increase the multiplexing capability of a dPCR system by detecting targets in multiple different fluorescent channels instead of only one as is classically done in dPCR.

Compared to existing methods for high-order multiplexing, the key advantage of the proposed color combination approach is the simplicity of data analysis. Here, a single threshold is required for each color to distinguish positive and negative droplets. The thresholding can be visualized on 1D plots and can be easily and quickly adjusted by the user. Lastly, the threshold values are robust to variations in the fluorescence values of positive droplets that occurs in practice.

With a read out in six fluorescent channels, we reliably detected up to 15 targets by combining two colors per target. Interestingly, combining two colors per target does not result in a major loss in sensitivity when the background concentrations of targets does not exceed a few hundreds of copies/μl. In addition, we showed that the sensitivity of the 1C-1T approach can be recovered by allocating one color to the abundant wild-type target and combining the other colors available to detect rare mutations.

Multiplexing scales quadratically with the number of colors available on the readout device when using color combination, in contrast with a linear scaling for all other high-order multiplexing approaches. With color combination and two colors per targets, 15 targets can be read out on a 6-color system, 21 targets on a 7-color system and 28 targets on an 8-color system.

The multiplexing capability can be increased even further with color combinations by labelling targets with only one color on top of labelling targets with two colors. For a 6-color system, this increases multiplexing capability to 21 (six targets are detected with one color per target on top of 15 targets with two colors per targets). However, the statistical framework to analyze the data is different from the description above and will be described elsewhere. Up to 20 targets could be also detected on a 6-color system by labelling each target with three colors.

Lastly, the price per point of a 15-plex assay with color combination can be significantly lower than NGS with reagent costs per assay and per sample typically below 1€. This competitive price makes color combination a viable approach to implement in routine testing of 15 to 30 targets in clinical diagnostics. This multiplexing capability will be highly beneficial in wide-ranging clinical applications such as liquid biopsy for cancer therapy selection or monitoring, as well as for infectious disease diagnostics or cell line and sample identification.

## Materials and Methods

### Reagents

Primers and probes used in the assays were synthesized by either Eurogentec (Seraing, Belgium) or IDT (Integrated DNA Technologies, Coralville, Iowa, USA) as indicated in the table in the Supp. Mat. with the indicated sequences and concentrations used. For the 15-plex assay, dsDNA gBlocks™ fragments (IDT) were used, corresponding sequences are indicated in the Supp. Mat. DNA concentrations used are indicated in the figures.

### dPCR

dPCR was performed using the Crystal Digital PCR™ technology from Stilla Technologies. First, primers, probes and targets are mixed at the indicated concentrations with 1X Buffer A (naica® PCR mix) and 4% of Buffer B (naica® PCR mix). The mix is loaded into Ruby chips and placed into the naica® Geode, a combined droplet generator and thermocycler, following the manufacturer’s instructions. For the 15-plex assay and 10+1-plex assay, the program used for amplification was 3min at 95°C for denaturation followed by 60 cycles of 15sec at 95°C and 3sec at 62°C.

### Data acquisition and analysis

Ruby chips were scanned using naica® Prism6 system with the Crystal Reader v4.0 (Stilla Technologies) software in 6 different channels for visualization of different fluorophores. Data was analyzed using Crystal Miner v4.0 (Stilla Technologies). For the analysis, a compensation matrix was applied to eliminate the fluorescence spillover assigned into a particular channel into different channels. Manual thresholding to determine positive and negative droplets in every color was set up. To identify the positive droplets corresponding to a particular target, population editor tool was used as described in the Supp. Mat. Calculation of target concentration and interval of confidence are determined automatically by Crystal Miner v4.0 according to Poisson distribution.

## Acknowledgements

We thank Cécile Jovelet for constructive discussions and advice regarding multiplex assay design; Philippe Rinaudo, Ward de Spiegelaere, Jo Vandesompele and Matthijs Vinck for constructive statistical discussions. This paper was typeset with the bioRxiv word template by @Chrelli: www.github.com/chrelli/bioRxiv-word-template.

## References

1. Salipante, S. J. & Jerome, K. R. Digital PCR — An Emerging Technology with Broad Applications in Microbiology. Clin. Chem. 66, 117–123 (2020).

2. Basu, A. S. Digital Assays Part I : Partitioning Statistics and Digital PCR. SLAS Technol. 22, 369–386 (2017).

3. Madic, J. et al. Three-color crystal digital PCR. Biomol. Detect. Quantif. 10, 34–46 (2016).

4. Madic, J. et al. EGFR C797S, EGFR T790M and EGFR sensitizing mutations in non-small cell lung cancer revealed by six-color crystal digital PCR. Oncotarget 9, 37393–37406 (2018).

5. Whale, A. S., Huggett, J. F. & Tzonev, S. Biomolecular Detection and Quantification Fundamentals of multiplexing with digital PCR. Biomol. Detect. Quantif. 10, 15–23 (2016).

6. Zhong, Q. et al. Multiplex digital PCR: breaking the one target per color barrier of quantitative PCR. Lab Chip 11, 20167–74 (2011).

7. Rowlands, V. et al. Optimisation of robust singleplex and multiplex droplet digital PCR assays for high confidence mutation detection in circulating tumour DNA. Sci. Rep. 9, 1–13 (2019).

8. Oscorbin, I. P. et al. Multiplex Droplet Digital PCR Assay for Detection of MET and HER2 Genes Amplification in Non-Small Cell Lung Cancer. Cancers (Basel). 14, 1–20 (2022).

9. Du, Y. et al. Development and evaluation of a multiplex droplet digital polymerase chain reaction method for simultaneous detection of five biothreat pathogens. Front. Microbiol. 13, 1–12 (2022).

10. Zhu, X. et al. Development of a multiplex droplet digital PCR assay for detection of enterovirus, parechovirus, herpes simplex virus 1 and 2 simultaneously for diagnosis of viral CNS infections. Virol. J. 19, 1–9 (2022).

11. Appay, R. et al. Multiplexed Droplet Digital PCR Assays for the Simultaneous Screening of Major Genetic Alterations in Tumors of the Central Nervous System. Front. Oncol. 10, 1–9 (2020).

12. Huang, Q. et al. Multicolor combinatorial probe Coding for real-time PCR. PLoS One 6, (2011).

13. Huang, Q., Hu, Q. & Li, Q. Identification of 8 foodborne pathogens by multicolor combinational probe coding technology in a single realtime PCR. Clin. Chem. 53, 1741–1748 (2007).

14. Liao, Y. et al. Combination of fluorescence color and melting temperature as a two-dimensional label for homogeneous multiplex PCR detection. Nucleic Acids Res. 41, (2013).

15. Marras, S. A. E., Tyagi, S., Antson, D. O. & Kramer, F. R. Color-coded molecular beacons for multiplex PCR screening assays. PLoS One 14, 1–12 (2019).

16. Quan, P. L., Sauzade, M. & Brouzes, E. DPCR: A technology review. Sensors (Switzerland) 18, (2018).

