## Supplementary information for "Robust higher-order multiplexing in digital PCR by color-combination"

### Supplementary Material for
### Robust higher-order multiplexing in digital PCR by color-combination

Authors: Irene Santos-Barriopedro, Sylvain Ursuegui, Etienne Fradet, Rémi Dangla

Stilla Technologies, 94800 Villejuif, France

#### SI#1. Calculations for CI & LOD

<u>Equation 1:</u>

Equation (1) first line states:

$$p_T = \frac{N_{Pos\,T}}{N_{tot}}$$

Using normal approximation (Wald interval), the confidence interval for $p_{T_i}$ is:

$$p_T \pm z_c \sqrt{\frac{p_T(1-p_T)}{N_{tot}}}$$

Defining: $p_{T_{min}} = p_T - z_c\sqrt{\frac{p_T(1-p_T)}{N_{tot}}}$ and $p_{T_{max}} = p_T + z_c\sqrt{\frac{p_T(1-p_T)}{N_{tot}}}$, and using equation (1) second line $C_T = -\frac{\ln(1-p_T)}{v}$, the confidence interval for $C_T$ is therefore defined by:

$$C_{T_{min}} = -\frac{\ln\left(1-p_{T_{min}}\right)}{v}; C_{T_{max}} = -\frac{\ln(1-p_{T_{max}})}{v}$$

For a 95% confidence interval, $z_c = 1.96$, and we end with equation (1) third line.

<u>Equation 3:</u>

Equation (2) first line states:

$$p_{T_i} = \frac{N_{Pos\,T_i}}{N_{All\,Neg} + N_{Pos\,T_i}}$$

Using normal approximation (Wald interval), the confidence interval for $p_{T_i}$ is:

$$p_{T_i} \pm z_c \sqrt{\frac{p_{T_i}(1-p_{T_i})}{N_{All\,Neg} + N_{Pos\,T_i}}}$$

Also,

$$N_{All\,Neg} + N_{Pos\,T_i} = N_{tot} - N_{Pos\,T_{j \neq i}}$$

with $N_{Pos\, T_{j\neq i}}$ the number of droplets containing other targets than $T_i$ (including co-encapsulation with $T_i$).

Using Poisson law:

$$N_{All\, Neg} + N_{Pos\, T_i} = N_{tot} - (1 - e^{-vC_{j\neq i}})N_{tot}$$
$$= e^{-vC_{j\neq i}}\, N_{tot}$$

with $C_{j\neq i} = \sum_{j\neq i} C_{T_j}$ the sum of the concentrations of the targets other than $T_i$, and $v$ the droplet volume

Therefore:

$$p_{T_i} \pm z_c\, e^{\frac{v}{2}C_{j\neq i}} \sqrt{\frac{p_{T_i}(1 - p_{T_i})}{N_{tot}}}$$

Defining: $p_{T_{i_{min}}} = p_{T_i} - z_c\, e^{\frac{v}{2}C_{j\neq i}} \sqrt{\frac{p_{T_i}(1-p_{T_i})}{N_{tot}}}$ and $p_{T_{i_{max}}} = p_{T_i} + z_c\, e^{\frac{v}{2}C_{j\neq i}} \sqrt{\frac{p_{T_i}(1-p_{T_i})}{N_{tot}}}$, and using equation (1) second line $C_{T_i} = -\frac{\ln(1-p_{T_i})}{v}$, the confidence interval for $C_{T_i}$ is therefore defined by :

$$C_{T_{i_{min}}} = -\frac{\ln\left(1-p_{T_{i_{min}}}\right)}{v} \text{ and } C_{T_{i_{max}}} = -\frac{\ln\left(1-p_{T_{i_{max}}}\right)}{v}$$

For a 95% confidence interval, $z_c = 1.96$, and we end with equation (3) first line.

The equation (3) second line is obtained by replacing $N_{All\, Neg} + N_{Pos\, T_i}$ by $e^{-vC_{j\neq i}}\, N_{tot}$ in the LOD formula.

Experimental calculations of LOD.

The directions to calculate the LOD in dPCR described before were followed (Stilla Tech Note). First, the calculation of the Limit Of Blank (LOB) is required in order to calculate the LOD to determine the false positive. The LOB of a target is the positive events that can be detected in a negative control with a confidence level of 95%. The LOD is the minimal concentration that can be detected and distinguished from the background. For the analysis of the LOB different replicates were analyzed: 38 samples (color combination 11-plex), 39 samples (single color 11-plex), 45 samples (color combination 15-plex) and 32 samples (single color 15-plex). After that, in order to calculate the LOD, samples with small concentration of the target (0.5, 0.8, 1.2, 1.5, 1.8 copies/µl) were analyzed. The number of samples used per assay were 37 samples (color combination 11-plex), 38 samples (single color 11-plex), 39 samples (color combination 15-plex) and 40 samples (single color 15-plex).

#### SI#2. Structure of the 15-plex proof-of-concept assay in color combination, primers & probes

| Name | Sequence | Fluorophore | Quencher | Concentration | Company |
|---|---|---|---|---|---|
| ESR1 530X F | CTGTACAGCATGAAGTGCAAG | | | 0.5 | Eurogentec |
| ESR1 530X R3 | GCATGTAGGCGGTGG | | | 0.5 | Eurogentec |
| ESR1 L536P FAM | TGCCCCCCTATGACCTGCT | FAM | /ZEN/ /Iowa Black FQ/ | 0.2 | IDT |
| ESR1 L536P YY | TGCCCCCCTATGACCTGCT | Yakima Yellow | /ZEN/ /Iowa Black FQ/ | 0.2 | IDT |
| PhiX174 F | TCTTTCCAAGCAACAGCAG | | | 0.5 | Eurogentec |
| PhiX174 R | AATACTGACCAGCCGTTTGA | | | 0.5 | Eurogentec |
| PhiX174 ATTO700 | TCCGAGATTATGCGCCAAATGC | ATTO700 | BHQ3 | 0.35 | Eurogentec |
| PhiX174 ROX | TCCGAGATTATGCGCCAAATGC | ROX | Iowa Black RQ | 0.35 | IDT |
| Lambda F | GCCATTGTTTCTCTGTGGAG | | | 0.5 | Eurogentec |
| Lambda R | TCACGGTTCAGTTGTTCACC | | | 0.5 | Eurogentec |
| Lambda YY | ACTGATTGCCCGTCTCCGCT | Yakima Yellow | /ZEN/ /Iowa Black FQ/ | 0.12 | IDT |
| Lambda ROX | ACTGATTGCCCGTCTCCGCT | ROX | Iowa Black RQ | 0.2 | IDT |
| pBR322 F2 | CGTTGATGCAATTTCTATGCG | | | 0.5 | Eurogentec |
| PBR322 R | TCGATAGTGGCTCCAAGTAG | | | 0.5 | Eurogentec |
| PBR322 FAM | CGCCCAGTCCTGCTCGCTTCG | FAM | /ZEN/ /Iowa Black FQ/ | 0.2 | IDT |
| PBR322 ATTO700 | CGCCCAGTCCTGCTCGCTTCG | ATTO700 | BHQ3 | 0.4 | Eurogentec |
| ALB F | TGAAACATACGTTCCCAAAGAGTTT | | | 0.5 | Eurogentec |
| ALB R | CTCTCCTTCTCAGAAAGTGTGCATAT | | | 0.5 | Eurogentec |
| ALB YY | TGCTGAAACATTCACCTTCCATGCA | Yakima Yellow | /ZEN/ /Iowa Black FQ/ | 0.1 | IDT |
| ALB Cy5 | TGCTGAAACATTCACCTTCCATGCA | Cy5 | BHQ2 | 0.3 | Eurogentec |
| pUC18 L1 F2 | CCTGCAGGTCGACTCTAG | | | 0.5 | Eurogentec |
| puC18MCSL1 R | TGAGCGGATAACAATTTCACA | | | 0.5 | Eurogentec |
| puc18 ATTO700 | CCCGGGTACCGAGCTCGAATT | ATTO700 | BHQ3 | 0.3 | Eurogentec |
| puc18 YY | CCCGGGTACCGAGCTCGAATT | Yakima Yellow | /ZEN/ /Iowa Black FQ/ | 0.15 | IDT |
| TP53 c.830-840 Fw | TCCTATCCTGAGTAGTGGTAATC | | | 0.5 | IDT |
| TP53 c.830-840 Rv | CCTTTCTTGCGGAGATTCTC | | | 0.5 | IDT |
| Mut-spe2 TP53 R282W- YY | TGCGCCAGTCTCTCCCAG | Yakima Yellow | /ZEN/ /Iowa Black FQ/ | 0.2 | IDT |
| Mut-spe2 TP53 R282W- Cy3 | TGCGCCAGTCTCTCCCAG | Cy3 | Iowa Black RQSp | 0.45 | IDT |
| TP53 R248 Fw | CACCATCCACTACAACTACATG | | | 0.5 | Eurogentec |
| TP53 R248 Rv | TCCTGACCTGGAGTCTTC | | | 0.5 | Eurogentec |

| | | | | | |
|---|---|---|---|---|---|
| TP53 WT3 R248 ROX | CATGAACCGGAGGCCCATC | ROX | Iowa Black RQSp | 0.5 | IDT |
| TP53 WT3 R248 Cy3 | CATGAACCGGAGGCCCATC | Cy3 | Iowa Black RQSp | 0.25 | IDT |
| MRM1 U1B | GTGGATAAGGTCATCACCA | | | 0.5 | IDT |
| MRM1 L1B | CAAGGTGCTTAGGAACTCG | | | 0.5 | IDT |
| MRM1_P2 ATTO700 | ACGTCCCTCATTCTCTATGTGCC | ATTO700 | BHQ3 | 0.25 | Eurogentec |
| MRM1_P2 Cy3 | ACGTCCCTCATTCTCTATGTGCC | Cy3 | Iowa Black RQSp | 0.2 | IDT |
| ESR1 E380Q F | TGGATTTGACCCTCCATGAT | | | 0.5 | Eurogentec |
| ESR1 E380Q R | CCAGACGAGACCAATCATCA | | | 0.5 | Eurogentec |
| ESR1 E380_WT- FAM | TT{C}TA{G}{A}ATGTGCCTGG | FAM | BHQ1 | 0.3 | Eurogentec |
| ESR1 E380_WT- ROX | TT{C}TA{G}{A}ATGTGCCTGG | ROX | BHQ2 | 0.15 | Eurogentec |
| DEL19-Fw | GTGAGAAAGTTAAAATTCCCG TC | | | 0.5 | IDT |
| DEL19-Rv | CCACACAGCAAAGCAGAAAC | | | 0.5 | IDT |
| DEL19-REF-P Cy5 | CACATCGAGGATTTCCTTGTTGGC | Cy5 | /TAO// Iowa Black RQSp/ | 0.35 | IDT |
| DEL19-REF-P Cy3 | CACATCGAGGATTTCCTTGTTGGC | Cy3 | Iowa Black RQSp | 0.15 | IDT |
| PIK3CA E453K Forward | GATTACACA{G}ACACTCTAGTATC | | | 0.5 | IDT |
| PIK3CA E453K Reverse | TGATCCAGTAACACCAATAGG | | | 0.5 | IDT |
| PIK3CA E453K Cy5 | CAA{A}T{C}TT{T}T{A}{A}{T}C{C}A{T}G | Cy5 | Iowa Black RQSp | 0.3 | IDT |
| PIK3CA E453K ROX | CAA{A}T{C}TT{T}T{A}{A}{T}C{C}A{T}G | ROX | BHQ2 | 0.15 | IDT |
| ERBB2 V777 Forward | GTCTCCCATACCCTCTCAG | | | 0.5 | IDT |
| ERBB2 V777 Reverse 2 | TGGATGTCAGGCAGATG | | | 0.5 | IDT |
| ERBB2 V777L Cy5 | AGCCCA{A}ACC{A}GCCAT | Cy5 | Iowa Black RQSp | 0.3 | IDT |
| ESR1 V422-F1 | TCATAGGAACCAGGGAAAATG | | | 0.5 | IDT |
| ESR1 V422-R1 | CGAGATGATGTAGCCAGC | | | 0.5 | IDT |
| ESR1 V422-del1 FAM | TGTAGAG{G}G{C}ATGGAGATC | FAM | Iowa Black FQ | 0.4 | IDT |
| ESR1 V422-del1 Cy5 | TGTAGAG{G}G{C}ATGGAGATC | Cy5 | Iowa Black RQSp | 0.15 | IDT |
| TSN Forward3 | TTTCTCTTTCCTGGTACAGTC | | | 0.5 | IDT |
| TSN Reverse2 | CCG GAA TCC AGC TCA TTG | | | 0.5 | IDT |
| TSN FAM | ACCCCTCCACATCTCCACCTT | FAM | BHQ1 | 0.55 | Eurogentec |
| TSN ATTO550 | ACCCCTCCACATCTCCACCTT | ATTO550 | BHQ2 | 0.08 | Eurogentec |

{} LNA: locked nucleic acid

#### SI#3. Structure of the comparable 15-plex using 1C-1T, primers & probes

| Name | Sequence | Fluorophore | Quencher | Concentration | Company |
|---|---|---|---|---|---|
| ESR1 530X F | CTGTACAGCATGAAGTGCAAG | | | 0.5 | Eurogentec |
| ESR1 530X R3 | GCATGTAGGCGGTGG | | | 0.5 | Eurogentec |
| ESR1 L536P FAM | TGCCCCCCTATGACCTGCT | FAM | /ZEN/ /Iowa Black FQ/ | 0.25 | IDT |
| PhiX174 F | TCTTTCCAAGCAACAGCAG | | | 0.5 | Eurogentec |
| PhiX174 R | AATACTGACCAGCCGTTTGA | | | 0.5 | Eurogentec |
| PhiX174 ROX | TCCGAGATTATGCGCCAAATGC | ROX | Iowa Black RQ | 0.25 | IDT |
| Lambda F | GCCATTGTTTCTCTGTGGAG | | | 0.5 | Eurogentec |
| Lambda R | TCACGGTTCAGTTGTTCACC | | | 0.5 | Eurogentec |
| Lambda ROX | ACTGATTGCCCGTCTCCGCT | ROX | Iowa Black RQ | 0.25 | IDT |
| pBR322 F2 | CGTTGATGCAATTTCTATGCG | | | 0.5 | Eurogentec |
| PBR322 R | TCGATAGTGGCTCCAAGTAG | | | 0.5 | Eurogentec |
| PBR322 FAM | CGCCCAGTCCTGCTCGCTTCG | FAM | /ZEN/ /Iowa Black FQ/ | 0.25 | IDT |
| ALB F | TGAAACATACGTTCCCAAAGAGTTT | | | 0.5 | Eurogentec |
| ALB R | CTCTCCTTCTCAGAAAGTGTGCATAT | | | 0.5 | Eurogentec |
| ALB Cy5 | TGCTGAAACATTCACCTTCCATGCA | Cy5 | BHQ2 | 0.25 | Eurogentec |
| pUC18 L1 F2 | CCTGCAGGTCGACTCTAG | | | 0.5 | Eurogentec |
| puC18MCSL1 R | TGAGCGGATAACAATTTCACA | | | 0.5 | Eurogentec |
| puc18 ATTO700 | CCCGGGTACCGAGCTCGAATT | ATTO700 | BHQ3 | 0.25 | Eurogentec |
| TP53 c.830-840 Fw | TCCTATCCTGAGTAGTGGTAATC | | | 0.5 | IDT |
| TP53 c.830-840 Rv | CCTTTCTTGCGGAGATTCTC | | | 0.5 | IDT |
| Mut-spe2 TP53 R282W- YY | TGCGCCAGTCTCTCCCAG | Yakima Yellow | /ZEN/ /Iowa Black FQ/ | 0.25 | IDT |
| TP53 R248 Fw | CACCATCCACTACAACTACATG | | | 0.5 | Eurogentec |
| TP53 R248 Rv | TCCTGACCTGGAGTCTTC | | | 0.5 | Eurogentec |
| TP53 WT3 R248 Cy3 | CATGAACCGGAGGCCCATC | Cy3 | Iowa Black RQSp | 0.25 | IDT |
| MRM1 U1B | GTGGATAAGGTCATCACCA | | | 0.5 | IDT |
| MRM1 L1B | CAAGGTGCTTAGGAACTCG | | | 0.5 | IDT |
| MRM1_P2 Cy3 | ACGTCCCTCATTCTCTATGTGCC | Cy3 | Iowa Black RQSp | 0.25 | IDT |
| ESR1 E380Q F | TGGATTTGACCCTCCATGAT | | | 0.5 | Eurogentec |
| ESR1 E380Q R | CCAGACGAGACCAATCATCA | | | 0.5 | Eurogentec |
| ESR1 E380_WT- ROX | TT{C}TA{G}{A}ATGTGCCTGG | ROX | BHQ2 | 0.25 | Eurogentec |
| DEL19-Fw | GTGAGAAAGTTAAAATTCCCG TC | | | 0.5 | IDT |
| DEL19-Rv | CCACACAGCAAAGCAGAAAC | | | 0.5 | IDT |

| | | | | | |
|---|---|---|---|---|---|
| DEL19-REF-P Cy3 | CACATCGAGGATTTCCTTGTTGGC | Cy3 | Iowa Black RQSp | 0.25 | IDT |
| PIK3CA E453K Forward | GATTACACA{G}ACACTCTAGTATC | | | 0.5 | IDT |
| PIK3CA E453K Reverse | TGATCCAGTAACACCAATAGG | | | 0.5 | IDT |
| PIK3CA E453K ROX | CAA{A}T{C}TT{T}T{A}{A}{T}C{C}A{T}G | ROX | BHQ2 | 0.25 | IDT |
| ERBB2 V777 Forward | GTCTCCCATACCCTCTCAG | | | 0.5 | IDT |
| ERBB2 V777 Reverse 2 | TGGATGTCAGGCAGATG | | | 0.5 | IDT |
| ERBB2 V777L ATTO700 | AGCCCA{A}ACC{A}GCCAT | ATTO700 | BHQ3 | 0.25 | Eurogentec |
| ESR1 V422-F1 | TCATAGGAACCAGGGAAAATG | | | 0.5 | IDT |
| ESR1 V422-R1 | CGAGATGATGTAGCCAGC | | | 0.5 | IDT |
| ESR1 V422-del1 FAM | TGTAGAG{G}G{C}ATGGAGATC | FAM | Iowa Black FQ | 0.25 | IDT |
| TSN Forward3 | TTTCTCTTTCCTGGTACAGTC | | | 0.5 | IDT |
| TSN Reverse2 | CCG GAA TCC AGC TCA TTG | | | 0.5 | IDT |
| TSN FAM | ACCCCTCCACATCTCCACCTT | FAM | BHQ1 | 0.25 | Eurogentec |

### SI#4. 2D plots of raw data obtained using the 15-plex POC assay (no truncation)

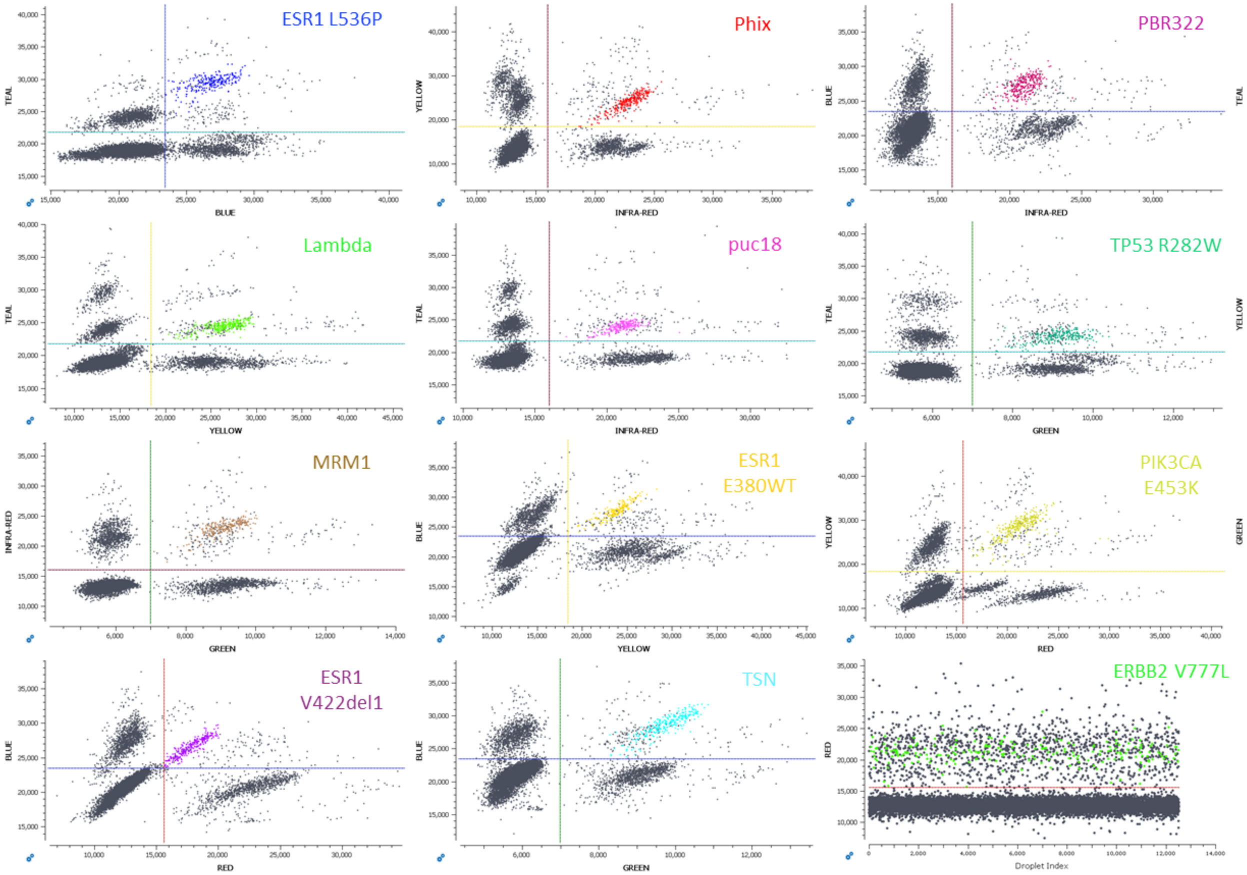


**2D-plots for 15-plex.** Diagrams of fluorescence intensity of two different channels. Each color corresponds to the droplets assigned to the different targets. Note the target 15 is detected only by the red channel

#### SI#5. Structure of the 11-plex proof-of-concept assay in color combination, primers & probes

| Name | Sequence | Fluorophore | Quencher | Concentration | Company |
|---|---|---|---|---|---|
| ESR1 530X F | CTGTACAGCATGAAGTGCAAG | | | 0.5 | Eurogentec |
| ESR1 530X R3 | GCATGTAGGCGGTGG | | | 0.5 | Eurogentec |
| ESR1 L536P FAM | TGCCCCCCTATGACCTGCT | FAM | /ZEN/ /Iowa Black FQ/ | 0.5 | IDT |
| ESR1 L536P YY | TGCCCCCCTATGACCTGCT | Yakima Yellow | /ZEN/ /Iowa Black FQ/ | 0.25 | IDT |
| PhiX174 F | TCTTTCCAAGCAACAGCAG | | | 0.5 | Eurogentec |
| PhiX174 R | AATACTGACCAGCCGTTTGA | | | 0.5 | Eurogentec |
| PhiX174 ATTO700 | TCCGAGATTATGCGCCAAATGC | ATTO700 | BHQ3 | 0.25 | Eurogentec |
| PhiX174 ROX | TCCGAGATTATGCGCCAAATGC | ROX | Iowa Black RQ | 0.25 | IDT |
| Lambda F | GCCATTGTTTCTCTGTGGAG | | | 0.5 | Eurogentec |
| Lambda R | TCACGGTTCAGTTGTTCACC | | | 0.5 | Eurogentec |
| Lambda YY | ACTGATTGCCCGTCTCCGCT | Yakima Yellow | /ZEN/ /Iowa Black FQ/ | 0.25 | IDT |
| Lambda ROX | ACTGATTGCCCGTCTCCGCT | ROX | Iowa Black RQ | 0.4 | IDT |
| pBR322 F2 | CGTTGATGCAATTTCTATGCG | | | 0.5 | Eurogentec |
| PBR322 R | TCGATAGTGGCTCCAAGTAG | | | 0.5 | Eurogentec |
| PBR322 FAM | CGCCCAGTCCTGCTCGCTTCG | FAM | /ZEN/ /Iowa Black FQ/ | 0.35 | IDT |
| PBR322 ATTO700 | CGCCCAGTCCTGCTCGCTTCG | ATTO700 | BHQ3 | 0.35 | Eurogentec |
| ALB F | TGAAACATACGTTCCCAAAGAGTTT | | | 0.5 | Eurogentec |
| ALB R | CTCTCCTTCTCAGAAAGTGTGCATAT | | | 0.5 | Eurogentec |
| ALB FAM | TGCTGAAACATTCACCTTCCATGCA | FAM | BHQ1 | 0.4 | Eurogentec |
| ALB Cy3 | TGCTGAAACATTCACCTTCCATGCA | Cy3 | /Iowa Black RQ/ | 0.2 | IDT |
| pUC18 L1 F2 | CCTGCAGGTCGACTCTAG | | | 0.5 | Eurogentec |
| puC18MCSL1 R | TGAGCGGATAACAATTTCACA | | | 0.5 | Eurogentec |
| puc18 ATTO700 | CCCGGGTACCGAGCTCGAATT | ATTO700 | BHQ3 | 0.25 | Eurogentec |
| puc18 YY | CCCGGGTACCGAGCTCGAATT | Yakima Yellow | /ZEN/ /Iowa Black FQ/ | 0.25 | IDT |
| TP53 c.830-840 Fw | TCCTATCCTGAGTAGTGGTAATC | | | 0.5 | IDT |
| TP53 c.830-840 Rv | CCTTTCTTGCGGAGATTCTC | | | 0.5 | IDT |
| Mut-spe2 TP53 R282W- YY | TGCGCCAGTCTCTCCCAG | Yakima Yellow | /ZEN/ /Iowa Black FQ/ | 0.25 | IDT |
| Mut-spe2 TP53 R282W- Cy3 | TGCGCCAGTCTCTCCCAG | Cy3 | Iowa Black RQSp | 0.5 | IDT |
| TP53 R248 Fw | CACCATCCACTACAACTACATG | | | 0.5 | Eurogentec |
| TP53 R248 Rv | TCCTGACCTGGAGTCTTC | | | 0.5 | Eurogentec |
| TP53 WT3 R248 ROX | CATGAACCGGAGGCCCATC | ROX | Iowa Black RQSp | 0.5 | IDT |
| TP53 WT3 R248 Cy3 | CATGAACCGGAGGCCCATC | Cy3 | Iowa Black RQSp | 0.25 | IDT |

| | | | | | |
|---|---|---|---|---|---|
| MRM1 U1B | GTGGATAAGGTCATCACCA | | | 0.5 | IDT |
| MRM1 L1B | CAAGGTGCTTAGGAACTCG | | | 0.5 | IDT |
| MRM1_P2 ATTO700 | ACGTCCCTCATTCTCTATGTGCC | ATTO700 | BHQ3 | 0.25 | Eurogentec |
| MRM1_P2 Cy3 | ACGTCCCTCATTCTCTATGTGCC | Cy3 | Iowa Black RQSp | 0.25 | IDT |
| ESR1 E380Q F | TGGATTTGACCCTCCATGAT | | | 0.5 | Eurogentec |
| ESR1 E380Q R | CCAGACGAGACCAATCATCA | | | 0.5 | Eurogentec |
| ESR1 E380_WT- FAM | TT{C}TA{G}{A}ATGTGCCTGG | FAM | BHQ1 | 0.25 | Eurogentec |
| ESR1 E380_WT- ROX | TT{C}TA{G}{A}ATGTGCCTGG | ROX | BHQ2 | 0.25 | Eurogentec |
| PIK3CA E453K Forward | GATTACACA{G}ACACTCTAGTATC | | | 0.5 | IDT |
| PIK3CA E453K Reverse | TGATCCAGTAACACCAATAGG | | | 0.5 | IDT |
| PIK3CA E453K Cy5 | CAA{A}T{C}TT{T}T{A}{A}{T}C{C}A{T}G | Cy5 | Iowa Black RQSp | 0.2 | IDT |

{} LNA: locked nucleic acid

#### SI#6. Structure of the comparable 11-plex assay using 1C-1T, primers & probes

| Name | Sequence | Fluorophore | Quencher | Concentration | Company |
|---|---|---|---|---|---|
| ESR1 530X F | CTGTACAGCATGAAGTGCAAG | | | 0.5 | Eurogentec |
| ESR1 530X R3 | GCATGTAGGCGGTGG | | | 0.5 | Eurogentec |
| ESR1 L536P FAM | TGCCCCCCTATGACCTGCT | FAM | /ZEN/ /Iowa Black FQ/ | 0.25 | IDT |
| PhiX174 F | TCTTTCCAAGCAACAGCAG | | | 0.5 | Eurogentec |
| PhiX174 R | AATACTGACCAGCCGTTTGA | | | 0.5 | Eurogentec |
| PhiX174 ROX | TCCGAGATTATGCGCCAAATGC | ROX | Iowa Black RQ | 0.25 | IDT |
| Lambda F | GCCATTGTTTCTCTGTGGAG | | | 0.5 | Eurogentec |
| Lambda R | TCACGGTTCAGTTGTTCACC | | | 0.5 | Eurogentec |
| Lambda ROX | ACTGATTGCCCGTCTCCGCT | ROX | Iowa Black RQ | 0.25 | IDT |
| pBR322 F2 | CGTTGATGCAATTTCTATGCG | | | 0.5 | Eurogentec |
| PBR322 R | TCGATAGTGGCTCCAAGTAG | | | 0.5 | Eurogentec |
| PBR322 ATTO700 | CGCCCAGTCCTGCTCGCTTCG | ATTO700 | BHQ3 | 0.25 | Eurogentec |
| ALB F | TGAAACATACGTTCCCAAAGAGTTT | | | 0.5 | Eurogentec |
| ALB R | CTCTCCTTCTCAGAAAGTGTGCATAT | | | 0.5 | Eurogentec |
| ALB FAM | TGCTGAAACATTCACCTTCCATGCA | FAM | BHQ1 | 0.25 | Eurogentec |
| pUC18 L1 F2 | CCTGCAGGTCGACTCTAG | | | 0.5 | Eurogentec |
| puC18MCSL1 R | TGAGCGGATAACAATTTCACA | | | 0.5 | Eurogentec |
| puc18 ATTO700 | CCCGGGTACCGAGCTCGAATT | ATTO700 | BHQ3 | 0.25 | Eurogentec |
| TP53 c.830-840 Fw | TCCTATCCTGAGTAGTGGTAATC | | | 0.5 | IDT |

| | | | | | |
|---|---|---|---|---|---|
| TP53 c.830-840 Rv | CCTTTCTTGCGGAGATTCTC | | | 0.5 | IDT |
| Mut-spe2 TP53 R282W- YY | TGCGCCAGTCTCTCCCAG | Yakima Yellow | /ZEN/ /Iowa Black FQ/ | 0.25 | IDT |
| TP53 R248 Fw | CACCATCCACTACAACTACATG | | | 0.5 | Eurogentec |
| TP53 R248 Rv | TCCTGACCTGGAGTCTTC | | | 0.5 | Eurogentec |
| TP53 WT3 R248 Cy3 | CATGAACCGGAGGCCCATC | Cy3 | Iowa Black RQSp | 0.25 | IDT |
| MRM1 U1B | GTGGATAAGGTCATCACCA | | | 0.5 | IDT |
| MRM1 L1B | CAAGGTGCTTAGGAACTCG | | | 0.5 | IDT |
| MRM1_P2 Cy3 | ACGTCCCTCATTCTCTATGTGCC | Cy3 | Iowa Black RQSp | 0.25 | IDT |
| ESR1 E380Q F | TGGATTTGACCCTCCATGAT | | | 0.5 | Eurogentec |
| ESR1 E380Q R | CCAGACGAGACCAATCATCA | | | 0.5 | Eurogentec |
| ESR1 E380_WT- ROX | TT{C}TA{G}{A}ATGTGCCTGG | ROX | BHQ2 | 0.25 | Eurogentec |
| PIK3CA E453K Forward | GATTACACA{G}ACACTCTAGTATC | | | 0.5 | IDT |
| PIK3CA E453K Reverse | TGATCCAGTAACACCAATAGG | | | 0.5 | IDT |
| PIK3CA E453K Cy5 | CAA{A}T{C}TT{T}T{A}{A}{T}C{C}A{T}G | Cy5 | Iowa Black RQSp | 0.25 | IDT |

{} LNA: locked nucleic acid

#### SI#7. gBlocks gene fragments used for the 15-plex and 11-plex experiments

| Name | Sequence |
|---|---|
| ESR1 L536P | gtagtcctttctgtgtcttcccacctacagTAACAAAGGCATGGAGCATCTGTACAGCATGAAGTGCA<br>AGAACGTGGTGCCCCCCTATGACCTGCTGCTGGAGATGCTGGACGCCCACCGCCTACATGC<br>GCCCACTAGCCGTGGGAGGGGCATCCGTGGAGGAGACGGAC |
| PhiX 174 | ACTGCTCGCGTTGCGTCTATTATGGAAAACACCAATCTTTCCAAGCAACAGCAGGTTTCCGA<br>GATTATGCGCCAAATGCTTACTCAAGCTCAAACGGCTGGTCAGTATTTTACCAATGACCAAATCAAAG |
| Phage Lambda | CAACTGGCGTAATCATGGCCCTTCGGGGCCATTGTTTCTCTGTGGAGGAGTCCATGACG<br>AAAGATGAACTGATTGCCCGTCTCCGCTCGCTGGGTGAACAACTGAACCGTGATGTCAGC<br>CTGACGGGGAC |
| puc18 | CCAAGCTTGCATGCCTGCAGGTCGACTCTAGAGGATCCCCGGGTACCGAGCTCGAATTCGT<br>AATCATGGTCATAGCTGTTTCCTGTGTGAAATTGTTATCCGCTCACAATTCCACACAACATAC<br>GAGCCG |
| pBR322 | TGCTGCTAGCGCTATATGCGTTGATGCAATTTCTATGCGCACCCGTTCTCGGAGCACT<br>GTCCGACCGCTTTGGCCGCCGCCCAGTCCTGCTCGCTTCGCTACTTGGAGCCACTATC<br>GACTACGCGATCAT |
| ALB | CGACCATGCTTTTCAGCTCTGGAAGTCGATGAAACATACGTTCCCAAAGAGTTTAAT<br>GCTGAAACATTCACCTTCCATGCAGATATATGCACACTTTCTGAGAAGGAGAGACAA<br>ATCAAGAAACAAACGT |
| ERBB2 V777L | AGGGCATAAGCTGTGTCACCAGCTGCACCGTGGATGTCAGGCAGATGCCCAGAAGGCGGG |

| | |
|---|---|
| | AGACATATGGGGAGCCCAAACCAGCCATCACGTATGCTTCCTGGGGACAAGGGTACGCTGA GAGGGTATGGGAGACCACACACCCCCAAACACCACACAGCCTCCCAACC |
| TSN | AACAAGCCCCTGTTTTCTCTTTCCTGGTACAGTCGAGGCTGTCTGTCAACAGCGTG ACTGCTGGAGACTACTCCCGACCCCTCCACATCTCCACCTTCATCAATGAGCTGGATTCCG GTTTTCGCCTTCTCAACCTGAAAAATGACTCCCTGAGGAAGCGCTACGACGGA |
| PIK3CA E453K | TGTTTGATTACACAGACACTCTAGTATCTGGAAAAATGGCTTTGAATCTTTGGCCAGTACCT CATGGATTAAAAGATTTGCTGAACCCTATTGGTGTTACTGGATCAAATCCAAATAAAGTAAG GTTTTTATTGTCATAAATTAGATATTTTTTATGGCAGTCAAACCGC |
| EGFRdel19 | ATCCCAGAAGGTGAGAAAGTTAAAATTCCCGTCGCTATCAAGGAATTAAGAGAAGCAACATC TCCGAAAGCCAACAAGGAAATCCTCGATgtgagtttctgctttgctgtgtgggggtccatgg |
| TP53 R248WT | CAAGCAGAGGCTGGGGCACAGCAGGCCAGTGTGCAGGGTGGCAAGTGGCTCCTGACCTGGAGT CTTCCAGTGTGATGATGGTGAGGATGGGCCTCCGGTTCATGCCGCCCATGCAGGAACTGTTACAC ATGTAGTTGTAGTGGATGGTGGTACAGTCAGAGCCAACCTAG |
| TP53 R282W | TAGTGCTCCCTGGGGGCAGCTCGTGGTGAGGCTCCCCTTTCTTGCGGAGATTCTCTTCCTCTGTGC GCCAGTCTCTCCCAGGACAGGCACAAACACGCACCTCAAAGCTGTTCCGTCCCAGTAGATTACCAC TACTCAGGATAGGAAAAGAGAAGCAAGAGGCAGTAAGG |
| ESR1 E380WT | TCAACTGGGCGAAGAGGGTGCCAGGCTTTGTGGATTTGACCCTCCATGATCAGGTCCACCTTCTAG AATGTGCCTGGCTAGAGATCCTGATGATTGGTCTCGTCTGGCGCTCCATGGAGCACCCAGGG |
| ESR1 V422del1 | tgcatgatctacgtgcgtcacatgcagtacatgttttcatagGAACCAGGGAAAATGTGTAGAGGGCATGGA GATCTTCGACATGCTGCTGGCTACATCATCTCGGTTCCGCATGATGAATCTGCAGGGAGAGG Agtcactagctcagattcagtagacgcgtgttg |
| MRM1 | GGGCTGTGCTGCGTTCCGCACACTTCCTCGGAGTGGATAAGGTCATCACCAGCCGGAGAAACAG GCACGGACGTCCCTCATTCTCTATGTGCCCCAACTTGGAGACGCAGCCGAGTTCCTAAGCACCTTG GCCCTTGGGTGATCCCTTAGCCAGACTTACCTGTCCCAGA |

### SI#8. Step by step instructions to set up Crystal Miner to analyze color combination assays

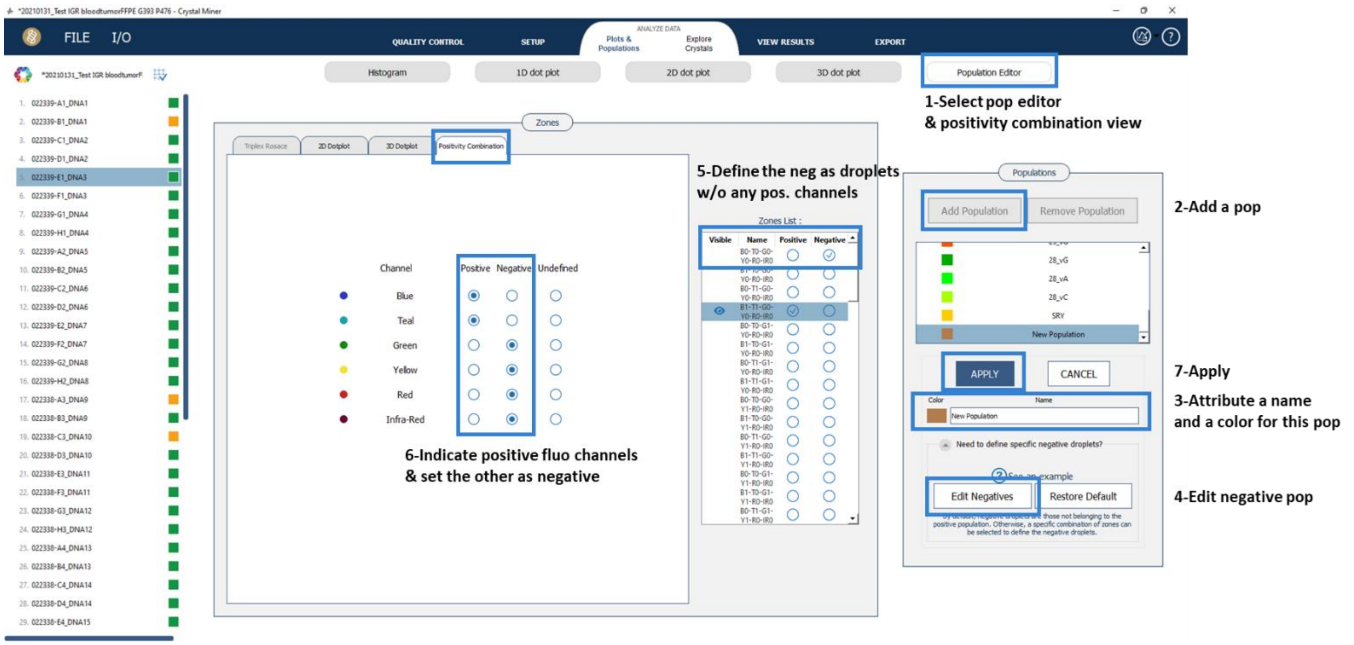


0. Place the thresholds in the 1D plot for the different channels/colors to properly separate negative droplets and positive droplets for each channel.
1. Click on “Population Editor” sub-tab in the “Plots & Populations” section of the software.
2. Select “Add population”
3. Write the name of the population, select the desired color for the population and click “OK”.
4. Click on “Need to define specific negative droplets?”; then click on “Edit Negatives”.
5. In the “Zone list”, select the first listed item (B0, T0, G0, Y0, R0, IR0) and only the first listed item as negative.
6. Click on the tab “Positivity Combination”. Select as positive the colors that are assigned for that population. The rest of the colors will be considered as negative.
7. Click on “Apply”.
8. Repeat the procedure for each population from the experiment.
9. To export the configuration, click on “I/O” and “Analysis Config”, select the “nca” file and save the analysis configuration as an “.nca” file. To import the configuration, select in “I/O” the option “Analysis Config” and choose the nca file.
10. In the "View Results” tab, the measurements for all defined populations will be displayed.
